# Pancreas-specific miR-216a regulates proliferation and endocrine and exocrine cell function *in vivo*

**DOI:** 10.1101/792655

**Authors:** Suheda Erener, Cara E. Ellis, Adam Ramzy, Maria M. Glavas, Shannon O’Dwyer, Sandra Pereira, Tom Wang, Janice Pang, Jennifer E. Bruin, Michael J. Riedel, Robert K. Baker, Travis D. Webber, Janel L. Kopp, Stephan Herzig, Timothy J. Kieffer

## Abstract

Pancreas is a vital organ composed of exocrine and endocrine cells that aid digestion of food and regulate blood glucose levels. Perturbations in the function of pancreatic cells leads to the development of life-burdening and/or threatening diseases such as diabetes and pancreatic cancer. Thus, it is critical to understand the molecular check-points that maintain normal pancreas physiology. MicroRNAs (miRNAs) are small non-coding RNAs involved in regulating gene expression in normal and diseased tissues. Several miRNAs have tissue-specific patterns consistent with crucial functions in many biological processes. Yet, there is limited knowledge about the role of pancreas-specific miRNAs in pancreatic pathologies. Here, we report that miR-216a is a conserved, pancreas-specific miRNA that is expressed in both endocrine and exocrine cells. Deletion of miR-216a in mice leads to reduced β-cell mass and a reduction in islet size under both chow and high-fat diet feeding conditions. We show that inhibition of miR-216a increases apoptosis and decreases cell proliferation in β- and exocrine cells. *Smad7* is upregulated in miR-216a deficient islets and cell cycle and proliferation are among the most significantly regulated biological processes in miR-216 knockout pancreata. Re-introduction of miR-216a in the pancreatic cancer line, PANC-1, increases cell migration more than 2-fold. *In vivo*, deletion of miR-216a in the pancreatic cancer prone mouse line *Kras*^*G12D*^;*Ptf1a*^*CreER*^ inhibits the propensity of pancreatic cancer precursor lesions. Our study identifies miR-216a as an important pancreas-specific miRNA which may have implications for both diabetes and pancreatic cancer.

## Introduction

The pancreas, an abdominal organ located behind the stomach, is composed of two major functional compartments. The exocrine pancreas, mainly comprised of acinar cells, aids digestion by secreting digestive enzymes amylase and lipase via pancreatic ducts into the duodenum while the endocrine pancreas, consisting of the islets of Langerhans, maintains normal glucose homeostasis. Islets are interspersed within the exocrine pancreas and consist of five different endocrine cells required for energy metabolism. The β-cells comprise 50-80% of the islets^1^ and are responsible for secreting insulin in response to changes in glucose levels. Perturbations in the function of endocrine and exocrine cells leads to the development of life-burdening and/or threatening diseases such as diabetes and pancreatic cancer. While diabetes is characterized by hyperglycemia resulting from defects in insulin secretion, insulin action, or both^2^, pancreatic ductal adenocarcinoma, a major form of pancreatic cancer in humans, results from the abnormal proliferation of the exocrine cells in response to oncogenic mutations^3^. To date many protein coding genes necessary for pancreas development and function have been identified, but the role of the majority of small RNAs is still unclear.

Micro-RNAs (miRNAs) are short (∼21-22 nt long) non-coding RNAs, which have emerged in the last two decades as buffers of signalling pathways to maintain normal tissue development and function^4^. Mature miRNAs function as evolutionarily conserved post-transcriptional gene regulators that mainly decrease the stability or inhibit translation of messenger RNAs (mRNAs) through binding to complementary sequences^5^. A single miRNA can impact the regulation of hundreds of genes with multiple targets^6^ within cellular networks that enable modulation of entire pathways in the context of an individual biological process^7^. Many miRNAs are conserved in sequence between distantly related organisms, suggesting that these molecules participate in essential processes. Recently, a study identified 3,707 novel mature miRNAs by analyzing 1,323 short RNA sequencing samples from 13 different human tissue types^8^, providing evidence that the repertoire of human miRNAs is far more extensive than that of public repositories and of what was previously anticipated^9^. Interestingly, many of the newly discovered miRNAs possess tissue specific patterns, akin to proteins. The functional impact of these miRNAs on various diseases remains to be investigated.

In the pancreas, conditional deletion of the miRNA processing endonuclease *Dicer* at the onset of pancreatic development (e9.5) using a Pdx1-Cre strain results in defects in all pancreatic lineages with dramatic reduction in the ventral pancreas as well as a reduction in the overall epithelial contribution to the dorsal pancreas at e18.5^10^. Postnatal *Dicer* ablation in the β-cells using the conditional *RIP2-Cre*^11,12^ or *Pdx1-CreER*^13^ strain impairs islet architecture, insulin secretion, and β-cell mass while deletion of *Dicer* in the acinar cells using the *Mist1-CreERT* mice^14^ promotes epithelial to mesenchymal transition accompanied by acinar to ductal metaplasia. Although previous findings demonstrate important roles for *Dicer* in the pancreas, *in vivo* studies investigating the role of individual miRNAs in pancreas development and endocrine and exocrine function are limited. Furthermore, the majority of investigated miRNAs do not have pancreas-specific expression.

Tissue-specific patterns of gene expression play fundamental roles in tissue development and function^15^. Although the majority of miRNAs are ubiquituosly expressed, some miRNAs exhibit tissue-specific^8^ or developmental-stage-specific expression patterns and contribute to maintaining normal tissue identity and function^16^. Furthermore, tissue-specific miRNAs are associated with various human diseases such as cardiovascular disease^17^ and cancer^18^. Therefore, it is critical to unravel the tissue specific regulatory networks to better understand the molecular mechanisms underlying diseases and identify new disease genes. In this study, we hypothesized that pancreas-enriched miRNAs have specific, critical roles for pancreas development and/or function and hence sought to identify miRNAs enriched in the pancreatic cells. Our results show that miR-216a is a pancreas-specific miRNA with critical functional roles in both endocrine and exocrine cells.

## Results

To identify the miRNAs that are enriched in pancreatic islets, we performed miRNA profiling from adult human islets and compared it to embryonic stem (ES) cells. We identified nine miRNAs that showed greater than 3-fold expression in human islets compared to ES cells **(Figure 1a)**. Out of the nine miRNAs identified, only miR-216a showed a pancreas-specific expression pattern **(Figure 1b)** while the eight other miRNAs were ubiquitously expressed in the analyzed tissues **(Suppl. Figure 1a)**. To examine whether miR-216a levels are changed during endocrine cell development, we differentiated human ES cells to pancreatic like cells., When comparing sequential timepoints, the single largest increase in levels of miR-216a occurred on day 14 **(Figure 1c)** of the differentiation protocol, which marks the generation of PDX1^+^/NKX6.1^+^ pancreatic endocrine progenitor cells^19^. miR-216a levels further increased during differentiation, reaching the highest levels at the final stage of differentiation (days 26-33), correlating with the presence of pancreatic endocrine cells. To further investigate miR-216a expression during development, we performed *in situ* hybridization with human fetal pancreatic tissue using DIG labelled miR-216a-LNA probes. There was a faint staining in the pancreatic tissue at the gestational time 8 weeks 4 days (8W 4d) (**Figure 1d**), during the time when insulin and glucagon double positive cells emerge^20^. However, we detected strong miR-216a staining in the branching pancreatic epithelium that harbor PDX1+ / NKX6.1+ pancreatic endocrine progenitor cells^20^. Analysis of human adult pancreas also revealed strong pancreatic miR-216a staining with further enrichment in pancreatic islets **(Figure 1d)**. To further investigate the specificity of miR-216a for pancreatic tissue, we performed *in situ* hybridization on kidney capsule grafts obtained from mice and rats implanted with pancreatic progenitor cells that had developed to endocrine cells^21^. There was strong reactivity for miR-216a in the grafts whereas neighboring kidney sections had no detectable staining **(Suppl. Figure 1b)**. We next examined the miR-216a sequence across diverse species and found it to be highly conserved **(Figure 1e)**, suggestive of a functional importance. A thorough tissue expression analysis from C57BL/6 mice showed miR-216a levels were specifically expressed in the pancreas with further enrichment in pancreatic islets **(Figure 1f)**, akin to our results with human tissues, suggesting a role for miR-216a miRNA in islet development and/or function.

**Figure 1.**
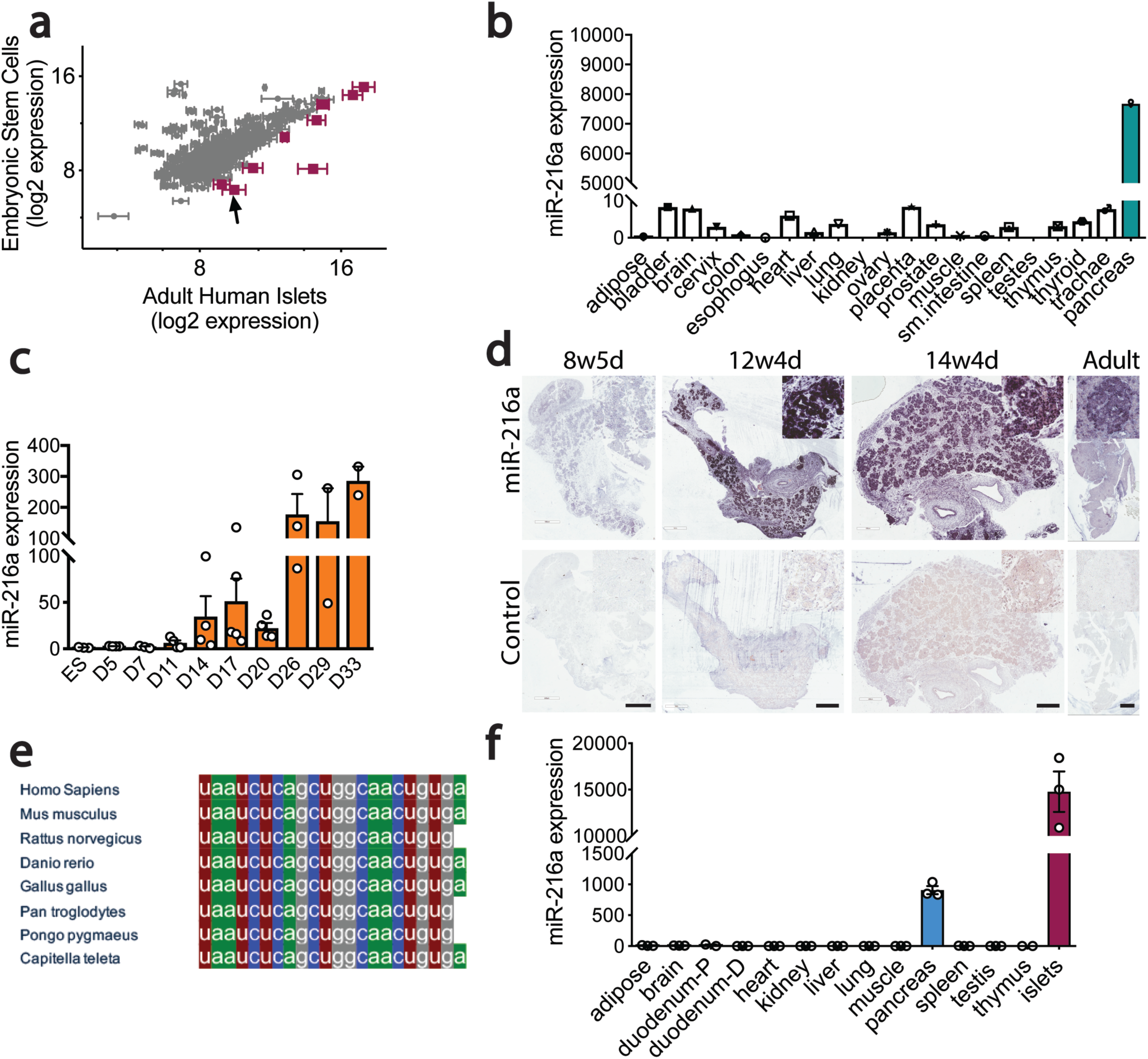
miR-216a is enriched in pancreatic islets and is highly conserved among various species. **a** miRNA profiling of adult human islets compared to human embryonic stem cells. miRNAs with greater than 3-fold increase in the islets are shown in red. Arrow points at miR-216a. n=3. Data represent mean log2 signals± SEM (from human islets). **b** Equal amounts of RNA from various human tissues (each a pool of 3 tissue donors) was reverse-transcribed and miR-216a expression was determined by qRT-PCR. Threshold cycle 33 (Ct= 33) was arbitrarily set as 1. **c** Human embryonic stem (ES) cells were differentiated to pancreatic endocrine cells for the indicated days and miR-216a expression was measured by qRT-PCR and expressed relative to levels in undifferentiated ES cells. **d** Fetal and adult human pancreata were probed with DIG-labeled miR-216a and scrambled control miRNA probes at the indicated gestational weeks. Purple color indicates presence of miRNA expression. Scale bar=l mm (12w4d, Adult), Scale bar= 500 µm (8w5d, 14w4d). Insets are enlarged 20x. **e** Comparison of mature miR-216a sequences in different species. **f** Same as in **(b)** except that the tissues were harvested from 8-week old CS7BL/6 male mice. n = 3. Individual data points are shown in c and f and data represent mean ± SEM.

To explore the role of miR-216a in pancreatic islet function *in vivo*, we generated miR-216a knock-out mice (miR-216a KO) in which the precursor sequence of miR-216a (pre-miR-216) was deleted by homologous recombination^22^. qRT-PCR analysis for miR-216a using total RNA from isolated islets of miR-216a KO mice revealed that miR-216a was not detected **(Suppl. Fig 2a)**. KO mice were viable and there were no significant differences in fasting body weight, blood glucose and insulin levels from newly weaned 3-4 weeks old miR-216a KO and littermate wild-type (WT) mice **(Suppl. Fig 2b-d)**. Pancreas weight and pancreatic cell size were unchanged **(Suppl. Fig 2e-f)**. We next monitored the miR-216a KO mice weekly for over 20 weeks for changes in body weight and blood glucose levels. miR-216a KO mice were comparable to WT littermate controls **(Figure 2a, b)**. Oral glucose tolerance tests performed at 10 and 15 weeks of age revealed no significant genotype differences in blood glucose **(Figure 2c, Suppl. Figure 2e)** or insulin levels **(Figure 2d)**. Similarly, miR-216a KO mice had normal insulin tolerance at 12 weeks of age and response to arginine injection at 18 weeks **(Figure 2e, Suppl. Figure 2f)**. We next assessed whether islets from miR-216a KO mice had altered insulin secretion *ex vivo*. Glucose (16.7 mM) stimulated insulin secretion from miR-216a KO islets was comparable to WT islets albeit insulin release was significantly lower at 2.8 mM glucose **(Figure 2f)**. Correspondingly, islets isolated from the miR-216a KO mice appeared generally smaller than the WT islets **(Figure 2g, h)**, and lacked the bigger islets with a trend towards an increased number of smaller islets **(Figure 2i)**. To analyze the effect of miR-216a KO on islet structure in more detail, we performed immunostaining with fixed pancreas tissue harvested from 21-week old adult mice. Immunostaining with insulin and glucagon antibodies demonstrated that β-cell mass was significantly reduced in the miR-216a KO mice while α-cell mass was unchanged **(Figure 3a-c)**. Islet circularity and the location of α-cells were unchanged **(Suppl. Fig 3a)**. Consistent with the isolated islet data, the area of islets determined by synaptophysin immunostaining was significantly smaller in miR-216a KO mice **(Figure 3a, d)** and miR-216a KO pancreatic sections harbored less of the larger islets **(Figure 3e)**. To examine whether the effects of miR-216a on β-cell mass and islet size was postnatally regulated, we performed the same analysis in newborn mice. Average islet size in one-day old miR-216a KO pups was comparable to WT littermates **(Figure 3f, g)**, with no statistical difference in various islet size groups **(Figure 3f, h)**. Interestingly, there was a trend towards an increased frequency for larger islets in the miR-216a KO pups **(Figure 3h)**. β-cell and α-cell area were not different as well as islet circularity and peripheral α-cells in one day old pups **(Suppl. Figure 3b-c)**.

**Figure 2.**
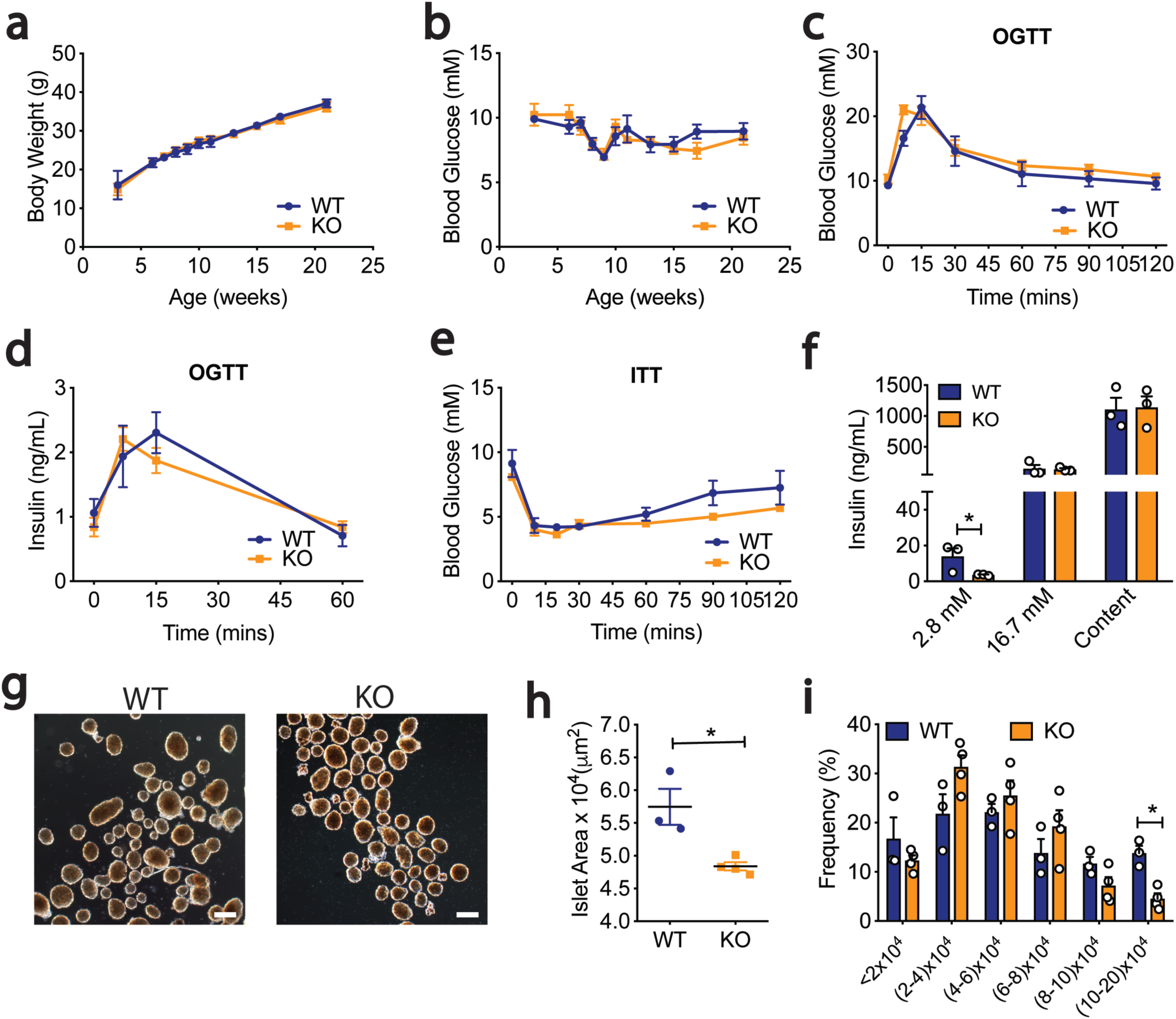
miR-216a knock-out (KO) mice have smaller islets than wild-type (WT) mice. WT and miR-216a KO male mice on regular chow diet were monitored for body weight **(a)** and blood glucose **(b)** for 21 weeks. n = 4-7. Oral glucose tolerance test (OGTT) performed in 10-week old male mice, with measurement of blood glucose levels **(c)** and plasma insulin concentration **(d). e** Insulin tolerance test (ITT) in 12-week old male mice injected with 1U/kg insulin at time = 0 and blood glucose levels determined at the indicated time points. **f** Insulin content and secretion from islets isolated from male WT and KO mice exposed to 2.8 mM and 16.7 mM glucose. n = 3. **g** Representative images of isolated islets from 10-week old male WT and KO mice. Scale bar = l00 µm. **h** Average size of isolated islets and the distribution of islet size **(i).** n = 3-4. A two-tailed Student’s t-test was performed to assess significance. *p<0.05 Individual data points are shown in **f**, **h**, and **i**. Data represent mean ± SEM.

**Figure 3.**
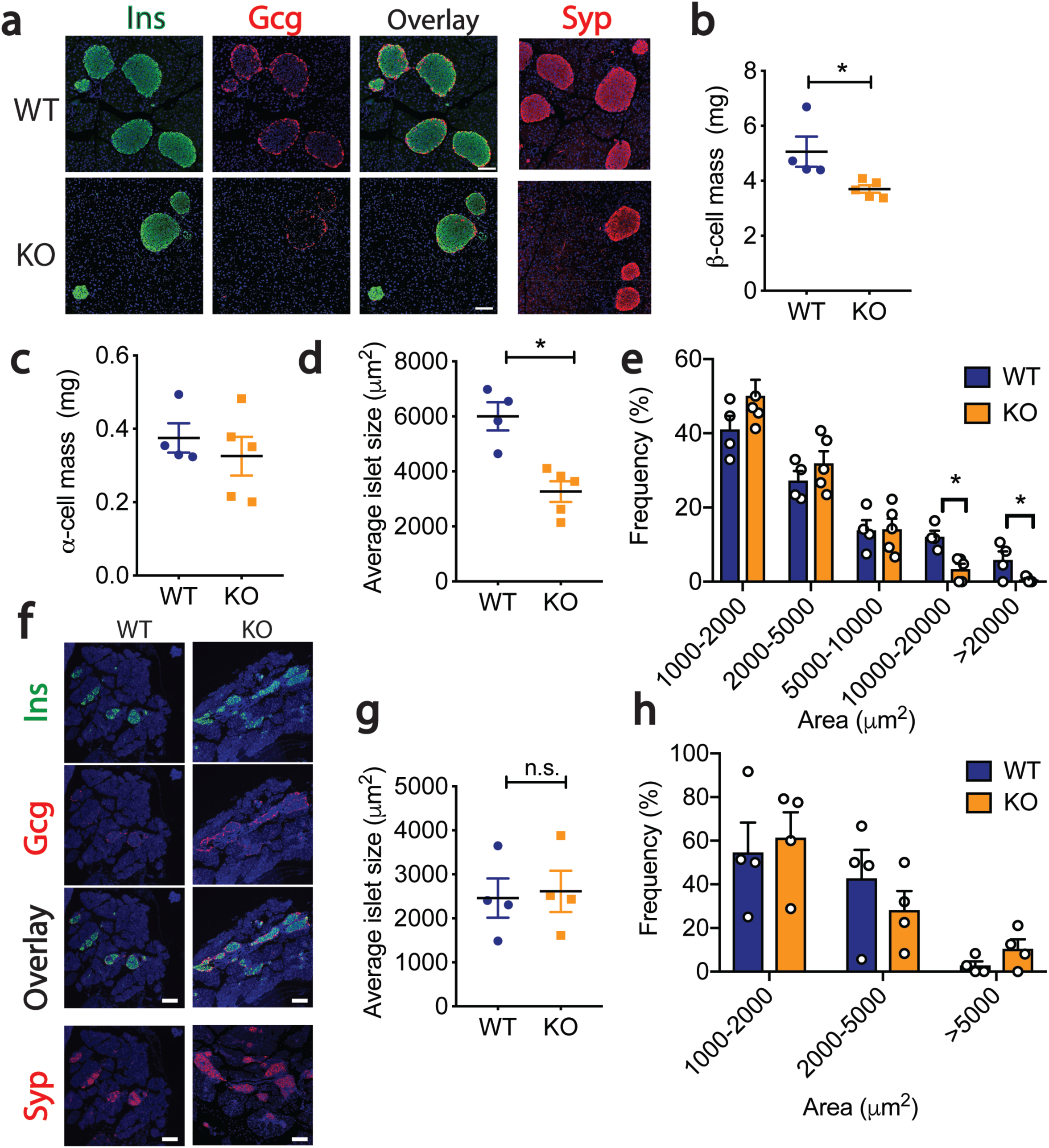
β-cell mass is reduced in miR-216a KO mice. **a-e** Pancreata from 21-week old male WT and KO mice were immunostained for insulin, glucagon and synaptophysin. **a** Representative images of insulin (Ins), glucagon (Gcg) and synaptophysin (Syp) immunostaining, along with an overlay of Ins and Gcg. Nuclei were identified with dapi (blue). Scale bars= 100 µm. **b** β-cell mass, **c** α-cell mass, **d** Average islet size, and **e** Islet size distribution. n = 4-5. **f-h** Pancreata from 1-day old male WT and KO mice were immunostained for insulin, glucagon and synaptophysin. **f** Representative images of insulin (Ins), glucagon (Gcg) and synaptophysin (Syp) immunostaining, along with an overlay of Ins and Gcg. Nuclei were identified with dapi (blue). Scale bar= 100 µm. **g** Average islet size, and **h** Islet size distribution. A two-tailed Student’s t-test was performed to assess significance. *p<0.05. Individual data points are shown in **(b-e, g-h).** Data represent mean ± SEM.

To explore the reasons for smaller islet size and reduced β-cell mass in miR-216a KO mice, we first examined the key hormones and transcription factors regulating endocrine cell identity. Expression of *Insulin, Glucagon, Pdx1*, and *Nkx6.1* were not changed in islets from miR-216a KO mice **(Suppl. Fig 4a)**. We next investigated whether miR-216a alters cell migration and proliferation. Regulation of islet size is complex and involves cellular processes such as fusion, fission, growth and migration^23^. Cell migration is critical for both islet formation and the movement of islets away from ducts^24,25^. We transfected pancreatic ductal adenocarcinoma PANC-1 cells that have very low miR-216a levels (not shown) and migration ability with control (ctrl) or miR-216 mimetics and performed a migration assay using transwell chambers. Quantification of the cells that traversed the boyden chambers demonstrated that miR-216a more than doubled the number of migrating cells compared to untransfected and control miRNA transfected wells **(Figure 4a)**. We next assessed the migration ability of miR-216a KO islet cells *ex vivo*. We coated cell culture plates with the matrix secreted by 804G cells and monitored the spreading of islets by light microscopy. Five days post-seeding, more WT islets spread compared to miR-216a KO islets while miR-216a KO islets had more defined borders **(Figure 4b)**.

**Figure 4.**
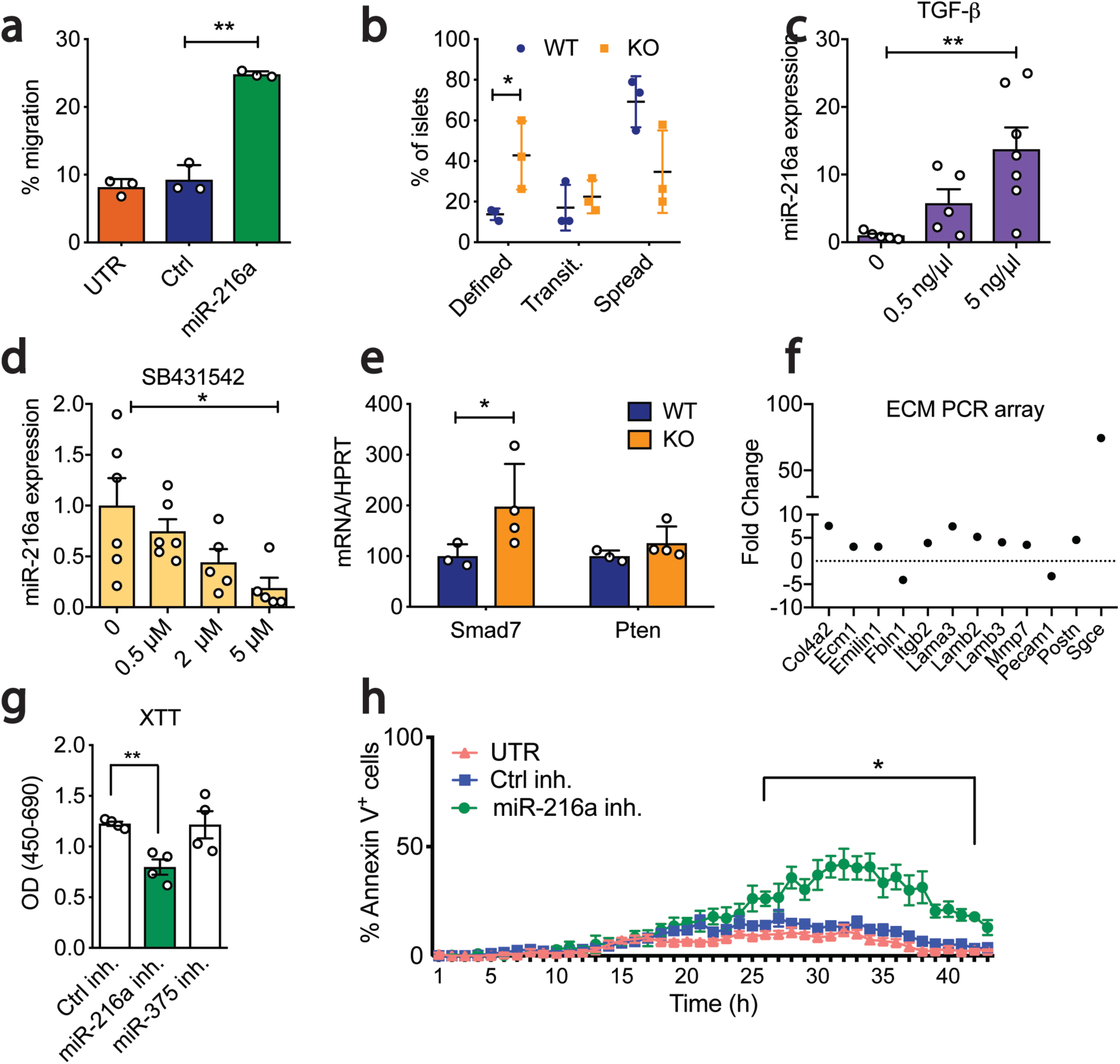
miR-216a regulates cell migration and apoptosis. **a** Percent of calcein stained PANC-1 cells migrating through a transwell following transfection with miR-216a or control (Ctrl) mimetics or untransfected (UTR). n = 3. **b** Percent of islets isolated from 10-week old male WT or KO mice that appeared as either having defined, transitionary (transit.) or spread boundaries three days after plating on collagen wells. n=3. **c-d** Human EndoC-βHl cells were treated with varying concentrations of **c** TGF-β or **d** TGF-β inhibitor SB431542 and miR-216a levels were quantified by qRT-PCR. n=6. **e** Islets-from 10-week old male WT and KO mice were isolated and expression of Smad7 and Pten was quantified with qRT-PCR. n = 3-4. **f** MIN6 cells were transfected with miR-216a and control mimetics and expression of 84 ECM related genes was quantified by qRT-PCR. Genes displaying > 2-fold difference are shown. Each value is the mean of 3 independent transfections. **g-h** INS1-E cells were transfected with the indicated miRNA inhibitors (inh.) or with a scrambled control miRNA inhibitor (Ctrl inh.) and cell viability was assessed by XTT (2,3-bis-(2-me thoxy-4-nitro-5-sulfophenyl)-2H-tetrazolium-5-carboxanilide) assay **(g)** or live cell imaging using Hoechst and Alexa647 annexinV **(h).** TNF-α, IFN-γ and IL-1β were added to media prior to imaging cells at 37°C and 5% CO2 in an lmageXpress Micro. n = 4. A two-tailed Student’s t-test **(a-e, g)** or two-way ANOVA with Bonferroni’s multiple comparison post-test (h) were performed to assess significance. *p<0.05, **p<0.01. Individual data points are shown in **(b-e, g-h).** Data represent mean± SEM.

TGF-β signalling regulates pancreatic epithelium branching and cell migration to form islet clusters^26^. It has been previously shown that miR-216a expression is regulated by TGF-β signalling and alters expression of Pten^27,28^ and Smad7^28^. To investigate the potential signalling pathways regulating miR-216a expression in the pancreas, we treated EndoC-βH1 cells, which have high endogenous miR-216a levels, with a TGF-β agonist TGF-β1 and the inhibitor SB431542 and measured miR-216a levels by qPCR. TGF-β1 treatment significantly increased miR-216 expression and inhibition of TGF signalling with SB431542 significantly decreased miR-216a levels **(Figure 4c-d)**. We next examined mRNA levels of potential miR-216a target genes in the pancreatic islets of WT and miR-216a KO mice. qRT-PCR analysis from islets of WT and miR-216a KO mice indicated that *Smad7* expression was significantly upregulated in the miR-216a KO islets whereas *Pten* levels were not changed **(Figure 4e)**. TGF-β signalling and Smad7 are known regulators of cell migration and their mode of action is thought to be via altering the expression of genes involved in regulating extracellular matrix composition^29^. We analyzed the levels of ECM genes in miR-216a transfected cells using an extracellular matrix (ECM) gene array. miR-216a altered the expression of 12 genes out of the 87 genes analyzed **(Figure 4f)**. Levels of genes involved in basement membrane such as *Col4a2, Ecm1, Fbln1, Lama3, Lamb2, Lamb3* were upregulated in the miR-216a transfected cells suggesting that TGF-β induced miR-216a can increase cell migration by altering the extracellular matrix integrity of basement membrane. We next investigated whether inhibition of miR-216a can affect cell proliferation and cell death by transfecting INS-1E β-cells with miR-216a and a control scrambled miRNA inhibitor. INS-1E β-cells have higher levels of miR-216a as compared to other rodent β-cell lines such as MIN6 (not shown) thus were chosen for miR-216a inhibition experiments. Inhibition of miR-216a significantly reduced cell proliferation compared to control miRNA inhibitors **(Figure 4g)**. Analysis of cell death using live cell imaging indicated that inhibition of miR-216a in the presence of cytotoxic factors (TNF-α, IFN-γ, IL-1β) increased the rate of apoptosis **(Figure 4h)** providing a possible mechanism for the smaller islet size.

Many miRNAs display their functional role under metabolic stress or upon cell insult^30^. We first explored whether miR-216a levels were altered in the islets of leptin knock-out (LepKO) rats, a model for metabolic overload with disturbed glucose and energy metabolism and larger islets^31^. Indeed miR-216a levels were significantly higher in the LepKO rat islets whereas levels of another well-studied islet enriched miRNA, miR-375, were unchanged **(Figure 5a)**. Although glucose metabolism in miR-216a KO mice was comparable to WT mice under chow-diet conditions, we next assessed whether reduced islet size and β-cell mass in miR-216a KO mice could result in impaired glucose homeostasis under high-fat diet conditions. miR-216a KO mice and the littermate WT controls were placed on 60% high-fat diet for eight weeks. Fasting body weight and blood glucose levels were comparable throughout the study **(Figure 5b-c)**. An oral glucose tolerance test performed 2 weeks after HFD feeding also revealed no significant differences in blood glucose levels **(Figure 5d)**, but miR-216a KO mice secreted less insulin in response to glucose administration **(Figure 5e)**. Similarly, repetition of the oral glucose tolerance test 6 weeks after HFD feeding showed no differences in blood glucose levels **(Figure 5f)**. However, miR-216a KO mice showed a trend towards reduced insulin secretion with significance at the basal time-point prior to oral glucose delivery **(Figure 5g)**. Insulin tolerance tests performed three and eight weeks after HFD feeding indicated similar insulin sensitivity of miR-216a KO and WT mice **(Figure 5h-i)**. Consistent with the reduced insulin secretion observed in the miR-216a KO mice, the animals had significantly reduced β-cell mass compared to controls **(Figure 5j,l**). In agreement with reduced β-cell mass, miR-216a KO mice had decreased circulating miR-375 levels **(Figure 5m)**, a miRNA that is highly expressed in the β-cells^32^ and is detected in the circulation^33,34^. We next measured plasma insulin and proinsulin levels to investigate whether the decreased β-cell mass was reflected in circulating insulin levels and whether β-cell work load was comparable between WT and miR-216a KO mice. Plasma insulin levels were comparable between WT and KO mice however, proinsulin levels were significantly decreased in the miR-216a KO mice **(Figure 5 n-o**), suggesting increased insulin processing in the KO β-cells to meet the metabolic demands. The degree of cell proliferation was comparable in the WT and miR-216a KO β-cells as assessed by PCNA immunostaining **(Figure 5 j,p**). To examine islet size, pancreata were immunostained for synaptophysin **(Figure 5 k**). Like mice on chow diet, miR-216a KO mice fed HFD also had significantly smaller islets **(Figure 5q)**, with increased frequency of smaller islets and decreased occurrence of larger islets **(Figure 5r)**. Overall, these data indicate that under metabolically stressed condition of HFD, miR-216a KO mice secrete lower insulin during a glucose challenge and have decreased islet size with reduced β-cell mass and increased markers of β-cell stress.

**Figure 5.**
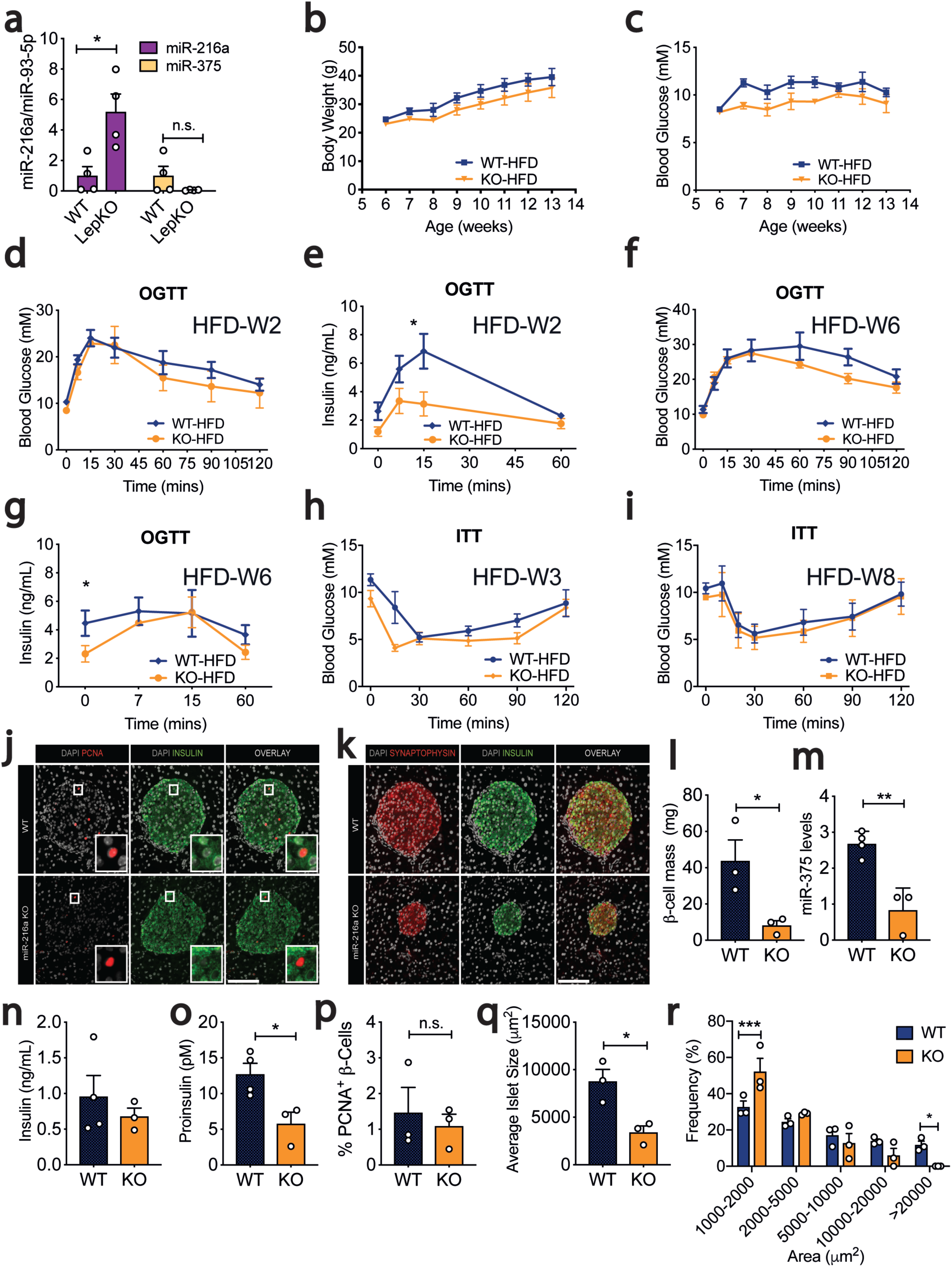
miR-216a KO mice have decreased β-cell mass and islet size on high-fat diet (HFD). **a** Islets were isolated from WT and Leptin knock-out male (LepKO) rats and expression of miRNAs was quantified by qRT-PCR and expressed relative to levels in WT cells. **b-r**WT and miR-216a KO male mice were fed with a 60% HFD for 8-weeks and weekly body weight **(b)** and fasted blood glucose levels **(c)** were measured. **d-g** Oral glucose tolerance tests (OGTT) were performed 2 weeks and 6 weeks post HFD, with measurement of blood glucose levels **(d, f)** and plasma insulin concentrations **(e, g). h**, **i** Blood glucose levels during insulin tolerance tests (ITT) performed 3 weeks **(h)** and 8 weeks (i) post HFD.**j-k** Pancreata from WT and miR-216a KO mice were fixed and stained with the indicated antibodies 8 weeks post HFD. Representative images are shown. Scale bars= 100 µm. Insets are enlarged 4x. **I**, **p, q** Quantifica tions of insulin **(I)**, PCNA **(p)** and synaptophysin **(q-r)** immunoreactivity. n = 3. **m** qRT-PCR analysis for miR-375 from plasma of WT and miR-216a KO mice, 8-weeks post HFD, n = 4. **n** Plasma insulin **o** Plasma proinsulin levels from WT and miR-216a KO mice, 8-weeks post HFD, n = 3-4.**q** Average islet size and **r** islet size distribution based on synapto physin immunostaining. A two-tailed Student’s t-test **(a, l-r)** or two-way ANOVA with Sidak’s multiple comparison post-test **(e, g)** were performed to assess significance. *p<0.05, *p<0.01, ***p<0.001. Individual data points are shown in **(a, l-r).** Data represent mean ± SEM.

Although we determined that *Smad7* expression was increased in the islets of miR-216a KO mice, we sought to perform a global analysis to further explore signalling pathways targeted by miR-216a in the whole pancreas. We performed RNA-sequencing from the pancreata of one day old WT and miR-216a KO mice. The RNA integrity number (RIN) obtained from the pancreata of miR-216a KO mice were all above 8 and suitable for RNA-sequencing (not shown). We examined the top 50 most abundant genes expressed in WT and miR-216a KO pancreata and observed the expected abundance for *Amylase, Trypsin* and *Insulin*, yet they were not significantly different between WT and miR-216a KO mice **(Figure 6a)**. Differential gene expression analysis from all the transcripts using an adjusted p-value < 0.05 revealed 409 genes were differentially expressed between WT and KO pancreata **(Suppl. table 1).** To identify more globally the affected biological processes in the miR-216a KO pancreata, we next performed a Gene Ontology analysis and found cell cycle to be most significantly altered **(Figure 6b)**. Similarly, KEGG pathway analysis indicated cell cycle, DNA replication and repair pathways were statistically different in the miR-216a KO versus WT pancreata **(Figure 6c)**. Next, we compared the expression of genes involved in cell cycle pathways between WT and miR-216a KO pancreata and observed that many cyclin dependent kinases (e.g: *Cdk4, Cdk1*) and cyclins (e.g: *Ccne1*) as well as DNA replication genes (e.g: *Pcna, Mcm2, Mcm5*) were decreased in the pancreas of miR-216a KO mice **(Figure 6d)**. This was consistent with our observation that inhibition of miR-216a decreases cell proliferation in INS-1E β-cells **(Figure 4g)** and the presence of reduced β-cell mass and islet size in the miR-216a KO mice. A map of the cell cycle pathway with genes significantly regulated in miR-216a KO pancreata is shown in **(Suppl. Fig 5)**. To investigate the effect of miR-216a on cell proliferation in cells with exocrine origin, we transfected human PANC-1 cells with the miR-216a mimetic and another miRNA, miR-217. Similar to INS1-1E β-cells, miR-216a transfection significantly increased the number of cells **(Figure 6e)**, and decreased the rate of apoptosis **(Figure 6f)**, thus confirming the role of miR-216a in regulating cell proliferation in another pancreatic cell type.

**Figure 6.**
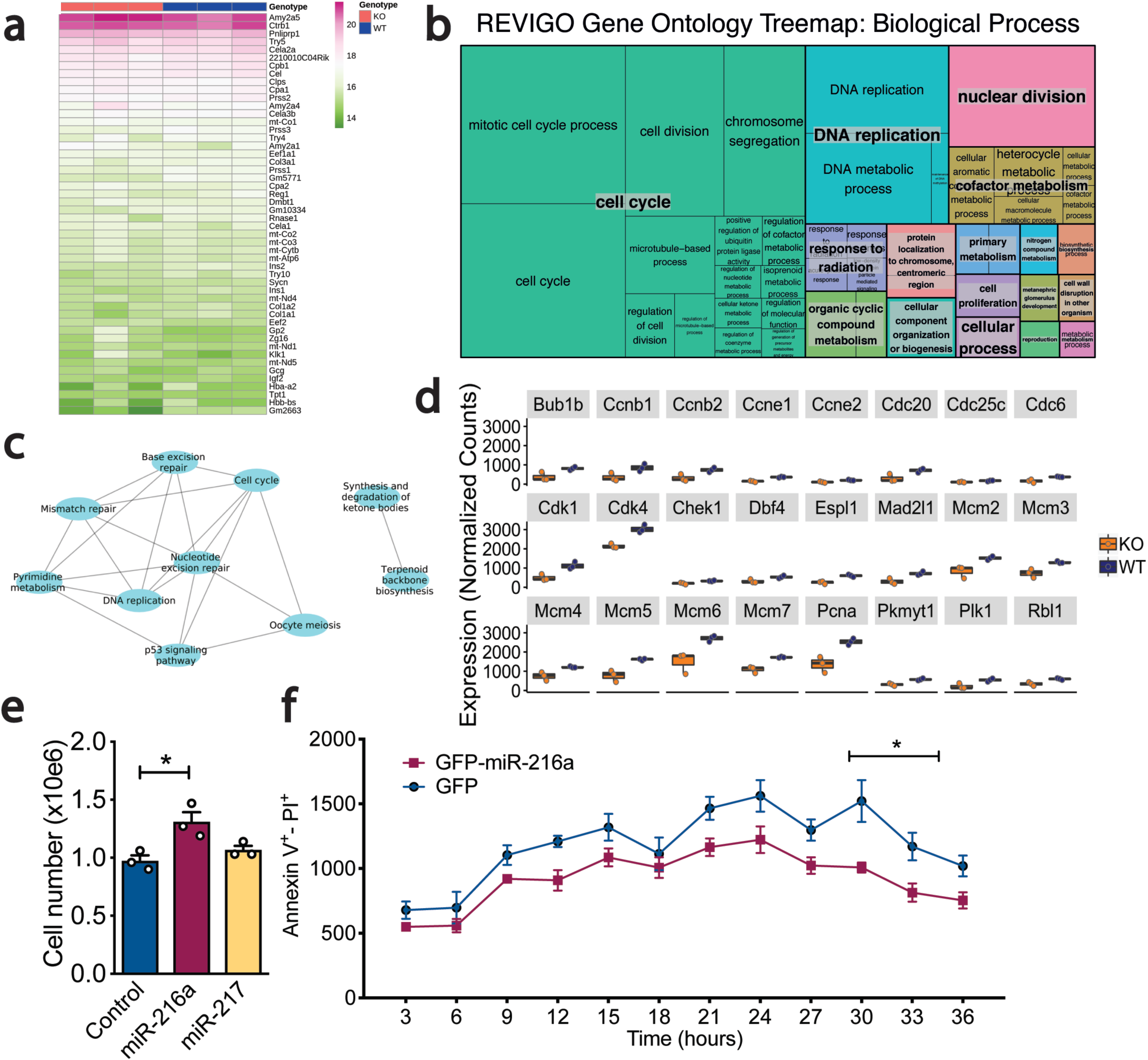
RNA-seq analysis of pancreata from miR-216a-KO mice. **a** RNA from the pancreata of 1-day old male WT and miR-216a KO mice was isolated and subjected to RNA sequencing and top 50 most abundant genes are shown by heat map. **b** A treemap plot, combining statistically significant GO terms in the biological processes category into similar term s. The size of the square increases with a decreasing p-value, and the colour of the square indicates the grouping of like terms (labelled in larger text with a grey background). Statistically significant terms were identified using hypergeometric tests with a false discovery rate of 0.1. **c** All statistically significant KEGG terms are shown in a network map, with nodes representing KEGG terms and edges connecting nodes representing differentially expressed genes in common between KEGG t erms. Statistically significant terms were identified using hypergeomet ric tests with a false discovery rate of 0.1. **d** Normalized gene expression data for key genes of interest. All genes shown are differentially expressed, with adjusted p-values < 0.05 (adjusted by the Benjamini Hochberg correct ion). **e-f** PANC-1 cells were transfected with the indicated miRNAs and 48-hours later TGF-β was added to cell culture media and cell number was counted **(e).** Individual data points are shown. n = 3. **f** Cell death was assessed with live cell imaging using Hoechst and Alexa647 annexinV. Cells were imaged at 37°C and 5% CO2 in an lmageXpress Micro. n = 5. A two-tailed Student’s t-test **(e)** or two-way ANOVA with Bonferroni’ s multiple comparison post-test **(f)** were performed to assess significance. *p<0.05. Data represent mean ± SEM, n= 3.

We next investigated whether the absence of miR-216a alters the progression of a pancreatic pathology related to cell cycle/proliferation in acinar cells. Oncogenic KRAS can induce pancreatic ductal adenocarcinoma (PDAC) precursor lesions from pancreatic acinar cells^35,36^. A number of studies have shown that PDAC develops from abnormally proliferating cells in the precursor lesions termed pancreatic intraepithelial neoplasia (PanIN)^37^, the most common precursor lesions observed in humans. A mutation in KRAS oncogene is currently considered as the initiating factor in pancreatic cancer^38^. Expression of constitutively active *Kras*^*G12D*^ allele in mice induces PanINs and after a significant latency period PDAC^39^. We crossed miR-216a KO mice with the pancreatic cancer prone *Kras*^*G12D*^;*Ptf1a*^*CreER*^ mice **(Figure 7a)** and analyzed the frequency of PanINs in the pancreata of offspring. As expected, H&E staining from the pancreata of *Kras*^*G12D*^;*Ptf1a*^*CreER*^;*miR-216a*^(*+/+)*^ mice revealed widespread lesions with columnar to cuboidal cells with varying degrees of cytological and architectural atypia **(Figure 7b)**. In contrast, the pancreata of mice lacking miR-216a had limited neoplastic lesions. Further histological analysis with Alcian blue staining confirmed the histological characteristic of high acidic mucin content of PanINs in the WT mice, whereas miR-216a KO pancreata had decreased PanIN frequency **(Figure 7c)**. In *Kras*^*G12D*^;*Ptf1a*^*CreER*^;*miR-216a*^(*-/-)*^ mice there was more than 2-fold reduction in the Alcian blue+ area compared to *Kras*^*G12D*^;*Ptf1a*^*CreER*^;*miR-216a*^(*+/+)*^ mice. **(Figure 7d)**. These data indicate that acinar cells from the miR-216a^(-/-)^ mice have less propensity than miR-216a^(+/+)^ mice to form PanINs and acinar to ductal metaplasia in response to oncogenic KRAS.

**Figure 7.**
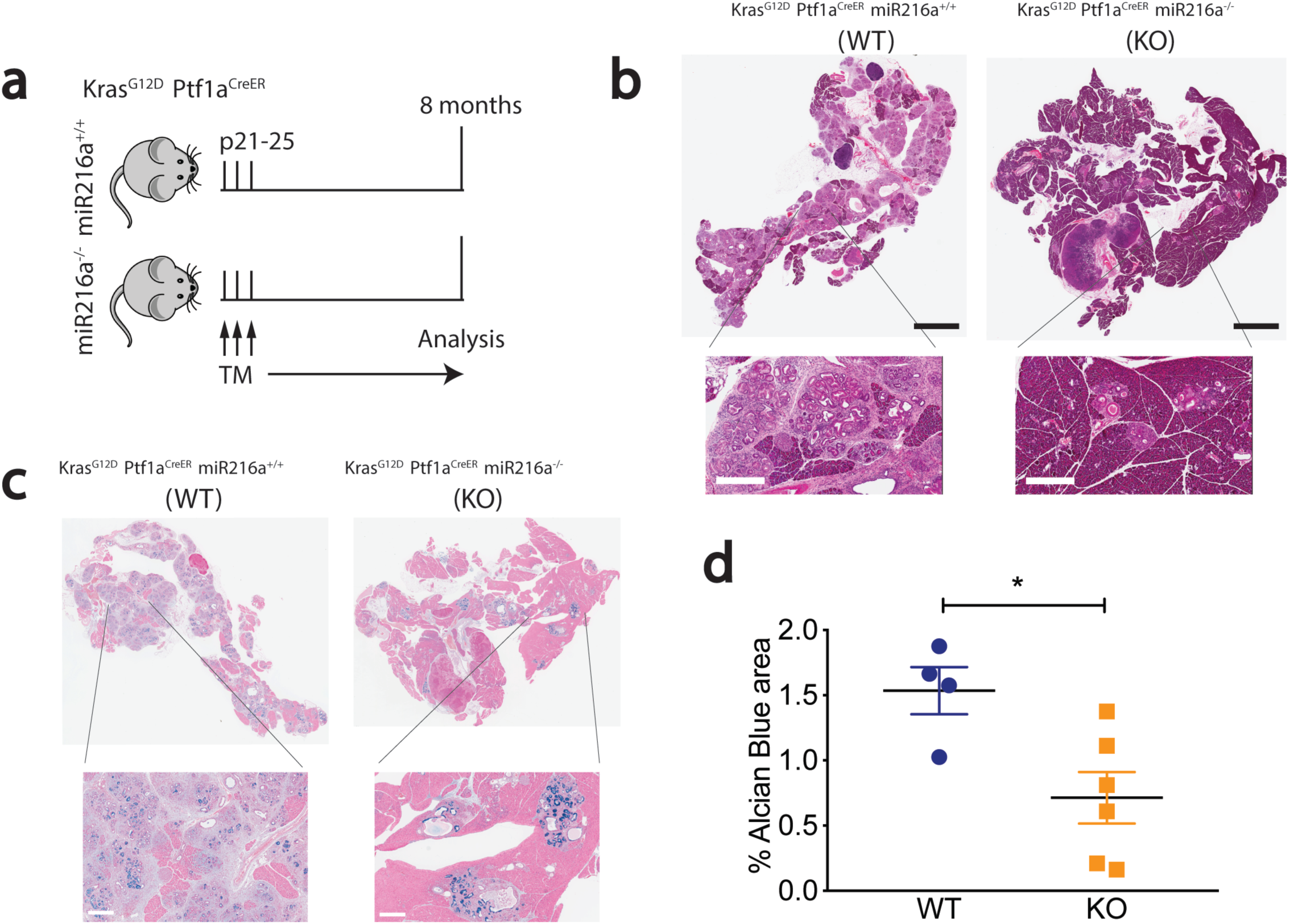
miR-216a-KO mice have lower incidence of pancreatic intraepithelial (PanIN) lesions. **a** Schematic showing the design of the study. Expression of KrasG12D was induced in exocrine cells by 3 tamoxifen injections on alternating days when the mice were 21-25 days old. Pancreata were harvested 8 months later. **b** Pancreatic H&E staining. Scale bars (black)= 3 mm, inset scale bars (white)= 400 µm. **c** Pancreatic alcian blue staining. Insets are 5x enlarged, inset scale bars= 400 µm. **d** Alcian blue positive area. Individual data points are shown. n = 4-6. A two-tailed Student’s t-test was performed to assess significance. Data represent mean ± SEM.*p<0.05.

## Discussion

We predicted that pancreas-enriched miRNAs likely have specific, critical roles for pancreas development and/or function and hence sought to identify miRNAs enriched in pancreatic cells. Our results show that miR-216 is a highly conserved pancreas-specific miRNA with roles in both endocrine and exocrine cell function. In the endocrine pancreas, we found that miR-216a regulates β-cell mass and islet size by increasing cell proliferation and migration. In the exocrine pancreas, miR-216a regulated cell cycle and proliferation and deletion of miR-216a decreased the formation of PanINs, implicating miR-216a in the progression of pancreatic pathologies such as diabetes and cancer.

miR-375 was the first reported islet “specific” miRNA^32^ with a role in maintaining normal α- and β-cell mass^45^. To our knowledge our study is the first to identify a pancreas-specific miRNA with functional roles in both endocrine and exocrine pancreas. Although we also observed an enrichment for miR-375 in human and mouse islets, it is also expressed at high levels in other tissues and non-pancreatic roles for this miRNA have been identified, such as in the regulation of gut mucosal immunity^46^ and gastric tumorigenesis^47^. Specificity of miR-216a for the pancreas suggests clinical opportunities to exploit this miRNA as a biomarker in pancreatic pathologies. Indeed, Goodwin et al. have shown that circulating miR-216 levels are increased in a mouse model of acute pancreatitis^48^. Szafranska et al. have shown that pancreatic tissue from PDAC patients have a large decrease in miR-216 expression^49^ and others have shown that miR-216 levels are lower in fecal specimens of patients with chronic pancreatitis and pancreatic cancer^50^. Further studies are required to evaluate the utility of miR-216a as serum biomarker of pancreatic pathologies and whether it can distinguish between different forms of pancreatic cancer and/or pancreatitis.

Using human and mouse tissue panels, fetal and adult human pancreas and human endocrine cell transplant models, we demonstrated that miR-216a is specific to pancreatic tissue. Therefore, a global miR-216a KO mouse was used to investigate the role of miR-216a in both endocrine and exocrine compartments *in vivo*. Islets isolated from the miR-216a KO mice were smaller and secreted less insulin than control islets. Similarly, analysis of pancreatic tissue confirmed the presence of smaller islets and reduced β-cell mass in the miR-216a KO adult mice, whether on normal or high fat diet. Interestingly, pancreata from the miR-216a KO newborn mice had a trend towards increased islet size. Islet growth is size-dependent during development, with preferential expansion of smaller islets and fission of large interconnected islet-like structures occurring most actively at approximately three weeks of age at the time of weaning^23^. As we observed a major difference in islet size in the miR-216a KO mice postnatally, our data supports a postnatal role for miR-216a in islet cell expansion and migration. Indeed, we show miR-216a increases cell migration and proliferation *in vitro*. Further studies are required to examine whether miR-216a can also impact cell fission.

Levels of miR-216a were regulated by TGF-β signalling and deletion of miR-216a led to an increase in *Smad7* levels in the miR-216a KO islets. This result is in alignment with published data showing that forced expression of miR-216a increases epithelial to mesenchymal transition (EMT) and migration with metastatic ability in epithelial hepatic cellular carcinoma (HCC) cells by targeting Pten and Smad7^28^. Although we did not observe any differences in Pten transcript or protein levels (not shown), we found an increase in *Smad7* levels in islets of miR-216a KO mice. We also observed changes in the expression of genes involved in EMT and migration in MIN6 and PANC-1 cells. Lack of Pten regulation by miR-216a in our study could be explained by differences in target gene selection under physiological vs. overexpression conditions such as the one used in the aforementioned studies. Alternatively, the selection of studied cell types (pancreas vs. HCC cells) could lead to differences in miRNA targets.

Analysis of RNA-seq results showed that cell cycle pathways were the most significantly enriched GO term in the miR-216a KO mice. Similarly, DNA replication, nuclear division, and cell proliferation were among the most altered biological processes. As miRNAs affect the expression of multiple genes, identifying signalling pathways targeted by miRNAs rather than minor changes in individual genes are more meaningful to interpret the role of miRNAs in the cellular and physiological context. By opting to isolate pancreatic RNA from one-day old pups, we focused on the earliest biological changes that occurred in the miR-216a KO mice postnatally, thereby revealing the potential causal mechanisms for the observed phenotype. Based on the expression of miR-216a in the acinar tissue as well as on the altered pathways in miR-216a KO pancreata, we hypothesized miR-216a may have a functional role in pancreatic cancer initiation. Our data show that deletion of miR-216 in the presence of oncogenic signals reduces incidence of neoplastic lesions in pancreatic cancer.

Cancer is characterized by uncontrolled proliferation resulting from aberrant activity of cell cycle proteins, thus many cell cycle regulators are important targets in cancer therapy^51^. A hallmark genetic event in PDAC is loss of the CDKN2A/2B tumor suppressor locus^52^ which encodes Cdk4/6 inhibitors that are particularly important for KRAS driven tumors such as PDAC^53,54^. Our RNA-seq analysis identified downregulation of many cell cycle and proliferation genes such as *Cdk1, Cdk4* and *Pcna* in the miR-216a KO mice and we found reduced PanIN formation in the presence of oncogenic KRAS, both of which suggest partial protection from pancreatic cancer initiation. TGF-β signalling pathway is another important and commonly deregulated signalling pathway in pancreatic carcinomas^55^. Activation of TGF-β leads to phosphorylation of Smad complexes to translocate to the nucleus and activate their target genes involved in cell cycle progression^55^. We showed that miR-216a levels are regulated by TGF-β signalling and miR-216a deletion increases levels of *Smad7*, which is a known inhibitor of cell cycle progression. Our results support a model whereby TGF-β induced miR-216 directly reduces *Smad7* levels and increases the expression of genes involved in cell cycle progression which in the presence of oncogenic mutations such as KRAS give rise to PanIN lesions. Whether there are other direct miR-216a targets mitigating this effect needs further evaluation. MiRNA based therapies are becoming more attractive due to their stability, sequence specificity and relative ease of miRNA synthesis^56^. Recently, engineered exosomes carrying short hairpin RNA specific to oncogenic Kras^G12D^, targeted oncogenic KRAS with an enhanced efficacy and suppressed cancer in multiple mouse models of pancreatic cancer^57^. It remains to be investigated whether inhibition of miR-216a either alone or in combination with KRAS can lead to a greater attenuation of tumor growth and an increase in overall survival in PDAC.

In summary, our results reveal for the first time a pancreas-specific miRNA (miR-216a) with physiological roles in both endocrine and exocrine cells, and show how dysregulation of miR-216a levels affect β-cell mass and pancreatic cancer initiation. miR-216a is upregulated by TGF-β signalling and pancreata of miR-216a KO mice harbor reduced size islets with reduced β-cell mass. miR-216a regulates the expression of genes involved in cell cycle progression and proliferation and deletion of miR-216a reduces the incidence of neoplastic lesions in KRAS mutant mice. Our findings offer insights into how islet size and β-cell mass are regulated by a pancreas-specific miRNA and offer potential intervention opportunities to slow β-cell death in diabetes. Furthermore, it opens up new possibilities to study the signalling pathways altered in PDAC patients. Finally, given the specificity of miR-216a for the pancreas, it provides biomarker opportunities to evaluate pancreatic disease status or progression.

## Materials and Methods

### Generation of MiR-216a Knockout Mice

The miR-216a KO ES cell line was generated by the Wellcome Trust Sanger Institute^22^ and the *miR-216a-/-* (miR-216a KO) mouse line was generated by the Centre for Phenogenomics in Toronto, Canada. Briefly, two frozen miR-216a KO ES cell (JM8A3 derived from C57BL/6N) clones (4H7 and 4H9) were purchased from the Mutant Mouse Resource and Research Center (USA) and were expanded and aggregated with diploid CD1 embryos. Coat color chimerism was scored and chimeras were bred with albino B6N (B6N-*Tyr*^*c-Brd*^/BrdCrCrl) females to test germline transmission. Successful germline transmission was confirmed with TaqMan qPCR of DNA prepared from tail clips, using the following primers in two separate reactions (KO and WT allele). The KO allele assay yields a 125 bp product. Forward: 5’AGT TCC TAT TCC GAA GTT CCT ATT C 3’. Reverse: 5’ AGA GGT TGA GGA CAG ACA GTA 3’. Probe (sense); TGG TCA TAG CTG TTT CCT GAA CAC CA. Cycling conditions: 98°C, 30s; 40 cycles of 98°C, 5s; 56°C, 10s. The WT allele assay yields a 135 bp product. Forward: 5’ GGC TAT GAG TTG GTT TA 3’. Reverse: 5’ GGA AAT TGC TCT GTT TAG 3’. Probe (antisense): CTG TGA GGA ATG ATA GGG AC. Cycling conditions: 95°C, 30s; 35 cycles of 95°C, 5s; 59°C, 10s. Reactions were performed using the iTaq Universal Probes Supermix (Bio-Rad #1725131, Hercules, California, USA) on the CFX96 Touch Thermal Cycler (Bio-Rad 1855195). Both probes were labelled with 6’-FAM.

M*iR-216a-/-* mice were crossed with WT C57BL/6 mice for at least 3 generations before use in experiments and maintained on a C57BL/6 background obtained from the University of British Columbia Animal Care Facility. In all experiments, aged-matched littermates of miR-216a KO mice were used as controls. Mice were fed a chow diet (2918, Research Diets) and were housed with a 12-h:12-h light-dark cycle with constant temperature and humidity and ad libitum access to food and water. Body weight and blood glucose were measured after a 4-hour morning fast. Blood was collected from the saphenous vein with heparin coated capillary tubes. All procedures with animals were approved by the University of British Columbia Animal Care Committee and carried out in accordance with the Canadian Council on Animal Care guidelines. For high-fat-diet (HFD) studies, C57BL/6 wild-type littermates and miR-216a KO mice were fed a 60% HFD diet (D12492i, Research Diets, Cedarlane, Burlington, Canada) starting at 6-8 weeks of age for 8 weeks. LSL-KrasG12D^58,59^ (Kras^G12D^) and Ptf1aCre^60^ have been described previously^35^ and were maintained on a mixed background. Leptin knock-out rats^31^ and kidney capsule grafts obtained from mice and rats implanted with pancreatic progenitor cells^21^ are described elsewhere.

### Metabolic Assessments

All metabolic analyses were performed in conscious mice that were restrained during blood sampling. Blood glucose values were determined with a OneTouch Ultra 2 glucometer (LifeScan, Inc., Burnaby, Canada) measured from saphenous vein blood. Fasting (4 hours) glucose and body weight were monitored weekly. For glucose tolerance tests with chow-diet or HFD fed mice, 6 hour fasted mice were given 2 g/kg glucose either by intraperitoneal injection or oral gavage, and blood was collected into heparin-coated capillary tubes at indicated times following glucose administration. For insulin tolerance tests, 4 hour fasted mice were injected intraperitoneally with 0.75 U/kg insulin (Novolin ge Toronto, Novo Nordisk Canada, Mississauga, Canada), with glucose measures at indicated times following glucose administration.

### Islet Isolation

Mouse islets were isolated as previously described^61^ using Type XI collagenase (1000 units/mL; #C7657, Sigma-Aldrich, St. Louis, MO). Following digestion and filtration, islets were picked with a pipette in three rounds to >95% purity. Islets were cultured overnight in RPMI-1640 (Sigma-Aldrich, St. Louis, MO) supplemented with 10% FBS (#F1051, Sigma-Aldrich, St. Louis, MO), 100 units/mL penicillin and 100 μg/mL streptomycin. For glucose stimulated insulin secretion assays, the next day after isolation 30 islets per mouse were placed in 24-well plates with KRBB buffer (129 mM NaCl, 4.8 mM KCl, 1.2 mM MgSO_4_, 1.2 mM KH_2_PO_4_, 2.5 mM CaCl_2_, 5.0 mM NaHCO_3_, 10 mM HEPES, 0.5 % BSA, pH: 7.4) supplemented with 2.8 mM Glucose (low-Glucose). After one hour of incubation at 37°C, media was discarded. Islets were then serially placed in low Glucose (2.8 mM Glucose) KRBB buffer, high Glucose (16.7 mM) KRBB buffer and 20 mM KCl for 1 hour at 37°C. At the end of each incubation media was collected, centrifuged and stored at - 20°C. Following the last media collection, islets were lysed in 0.18 M HCl/70% ethanol, homogenized, then incubated again at −20°C overnight. Following centrifugation, the aqueous solution was neutralized 1:2 with 1 M Tris, pH 7.5.

### Cell Culture

The INS-1 rat insulinoma cell line was cultured in RPMI-1640 containing 11.2 mM glucose and 2 mM L-glutamine. The medium was supplemented with 10% fetal bovine serum, 1 mM sodium pyruvate (#S8636, Sigma-Aldrich), 10 mM HEPES, 50 µM 2-mercaptoethanol (Sigma Aldrich), 100 U/mL penicillin (Sigma Aldrich), and 100 μg/mL streptomycin (Sigma Aldrich). PANC-1 cells were cultured in DMEM media (Life Technologies) containing 4 mM L-glutamine, 4500 mg/L glucose, 1 mM sodium pyruvate, and 1500 mg/L sodium bicarbonate supplemented with 10% fetal bovine serum, 100 U/ml penicillin, and 100 μg/ml streptomycin. EndoC-*β*H1 cells provided by (Drs. R Scharfmann, and P. Ravassard) were cultured on ECM-fibronectin–coated (1% and 2 μg/ml, respectively) (Sigma-Aldrich) culture wells and maintained in DMEM (Sigma Aldrich) that contained 5.6 mM glucose, 2% BSA fraction V (Roche Diagnostics), 50 μM 2-mercaptoethanol, 10 mM nicotinamide (Sigma-Aldrich), 5.5 μg/ml transferrin (Sigma-Aldrich), 6.7 ng/ml selenite (Sigma-Aldrich), 100 U/ml penicillin, and 100 μg/ml streptomycin. For transfections, cells were plated on 6-well plates (unless stated otherwise), and the next day 50-70% confluent cells were transfected with miR-216a (50 nM) and miRNA controls (50 nM) using RNAimax (Life Technologies) following manufacturer’s instructions. To control for tranfection effects, at least 3 wells were left untransfected. The protocol for differentiating CA1S human ES cells into pancreatic endocrine cells was described elsewhere^19^.

### Cloning

<u>miRNA-GFP expression vectors were</u> generated by initially creating a EGFP expression vector, pScamp EF1a-EGFP, which contains the 5’ end of the human EF1A gene (a genomic PCR amplicon containing 1407 bp of proximal promoter, exon 1, intron 1, and the UTR of exon 2), followed by the EGFP ORF and SV40 poly(A) signal, all cloned into pBluescript. This was subsequently modified into a “miRtron” expression vector, pScamp EF1a-premir-216a-EGFP, by removing 450 bp of the middle EF1A intron with restriction endonucleases BfuAI and XhoI, and replacing it with a compatible BsmBI-SalI restriction fragment of a 575-bp amplicon* consisting of the full hsa-premir-216a sequence, flanked on either side by roughly 200 bp of endogenous genomic sequences. Cells transfected with this plasmid produce a chimeric EF1A-EGFP mRNA that gets translated into a fluorescent protein, while the excised intron is processed into mature hsa-miR-216a.* miRtron hsa-premir-216a cloning primers (BsmBI and XhoI 5’ overhangs are underlined). Forward: 5’ gggcacacaCGTCTCG<u>ACGC</u>AGATTATACTTTTATGACATTACATGCAATATAGC 3’; Reverse: 5’ cacacaG<u>TCGA</u>CCCAAGTAGCACTGAAGGAGCG 3’.

### Pancreas Immunohistochemistry and Histology

Pancreata were fixed overnight in 4% paraformaldehyde (PFA) and then stored in 70% EtOH prior to paraffin embedding and sectioning (5 µm thickness; Wax-it Histology Services; Vancouver, BC). Immunofluorescent staining was performed as previously described^62^. Briefly, slides were deparaffinized in xylene and hydrated by graded ethanol washes. Slides were then washed in phosphate-buffered saline (PBS) followed by 10–15 min incubation in 10 mM sodium citrate/0.05% Tween 20, pH 6.0 at 95°C. Slides were incubated with DAKO Protein Block Serum-Free (Agilent Technologies, Inc, Santa Clara, CA) at room temperature for 10 minutes and then with the primary antibodies diluted in Dako Antibody Diluent (Agilent Technologies, Inc.) overnight at 4°C. The following primary antibodies were used: insulin (Cell Signalling, # 3014 at 1:200), glucagon (Sigma Aldrich, #G2654 at 1:1000), synaptophysin (Monosan, #MON9013 at 1:10), and PCNA (BD Transduction, #610665 at 1:100). The next day slides were serially washed with PBS and were incubated in appropriate Alexa Fluor 488 or 594 secondary antibodies (Thermo Fisher Scientific, Waltham, MA) diluted in Dako Antibody Diluent. Following serial PBS washes, slides were coverslipped with VECTASHIELD HardSet Mounting Medium with DAPI (Vector Laboratories, Burlingame, CA). Images were captured using a ImageXpress Micro™ Imaging System and analyzed using MetaXpress 5.3.0.5 Software (Molecular Devices Corporation, Sunnyvale, CA, USA). β-cell mass was analyzed in three pancreas sections per mouse, at least 200 µm apart, on insulin-labeled sections. Islet size was quantified as the synaptophysin-positive area per islet. For the α-cell analysis, glucagon-immunoreactive cells were counted using the ImageJ Cell Counter tool. Only islets with visible DAPI stained nuclei and at least 40 cells and 10 or more non-β cells were used to determine periphery or core localization of α-cells. α-cells were considered to be in the periphery of the islet if they were within 2 outermost cell layers of the islet or the core if they were deeper than the 2 outermost cell layers of the islet. For the islet circularity analysis, islets stained with synaptophysin and DAPI were used. Only islets with at least 40 cells and 10 or more non-β cells were used to determine islet circularity. All images were converted to 8-bit and non-islet nuclei were erased using ImageJ outline and clear function. Islet circularity was determined using Analyze Particles with threshold set until the entire area of islet was filled. H&E staining and Alcian blue staining on pancreatic sections were performed by Wax-it Histology Services Inc. (Vancouver, Canada).

### RNA Isolation and Quantitative RT-PCR

Tissues were homogenized with an Ultra-Turrax and total RNA was isolated using the miRCURY RNA isolation kit (Exiqon, U.S.A) as per the manufacturer’s instructions including a DNAse (Life Technologies) treatment step. mRNA was reverse transcribed with iScript cDNA Synthesis Kit (Bio-Rad Laboratories, Hercules, CA) and quantitative RT-PCR was performed using Ssofast EvaGreen Supermix (Bio-Rad). Primer sequences to amplify *Insulin, Pdx1, Nkx6.1*, and *Hprt* transcripts were described previously^63^. Primer sequences for *Glucagon* were Forward: 5’ CGCAGGCACGCTGATG 3’ Reverse: 5’ ACGAGATGTTGTGAAGATGGTTG 3’. A human RNA panel was purchased from ThermoFisher Scientific (#AM6000). miRNA was reverse transcribed with a Universal cDNA synthesis kit (Exiqon, U.S.A) and qRT-PCR was performed using SYBR Green master mix (Exiqon, U.S.A) with LNA-based miRNA primers (Exiqon, U.S.A). Relative values were calculated by the quantified by the 2^−ΔΔCT^ method. RNA from the pancreata of 1-day old mice was isolated using a protocol specific for pancreatic RNA isolations^64^.

### *In situ* Hybridization

Human fetal pancreata were kindly provided by Dr. Renian Wang, University of Western Ontorio, London, Canada and were described elsewhere^20^. Human adult pancreas was provided by the Irving K. Barber Human Islet Isolation Laboratory (Vancouver, British Columbia) with consent to use for research purposes. All samples were fixed overnight in 4% PFA, embedded in paraffin and sectioned (5 µm thickness; Wax-it Histology Services; Vancouver, BC). Slides were de-paraffinized with three consecutive xylene washes, rehydrated with graded ethanol washes (100% x 3, 95%, 70%, each 5 minutes), and washed in diethylpyrocarbonate (DEPC)-treated water. *In situ* hybridization was carried out with a IsHyb *In Situ* Hybridization kit (Biochain, San Francisco, California). Sections were fixed with 4% PFA for 20 minutes, washed twice with DEPC-treated PBS, and treated with 10 µg/mL Proteinase K (Sigma Aldrich) at 37°C for 15 minutes. Slides were washed with DEPC-treated PBS, fixed again with PFA for 15 minutes, and washed with DEPC-treated water. Sections were incubated for 4 hours at 50°C in pre-hybridization solution (Biochain). Afterwards, pancreatic sections were incubated with DIG-labeled hsa-miR-216a miRCURY LNA detection probe (#38495-15, Exiqon) or control scrambled miRNA probe (#99004-15, Exiqon) at 0.25 ng/µL in hybridization solution (Biochain) for 14 hours at 45°C. Then, slides were washed in SSC buffer (Biochain) as follows: 2X SSC buffer, 2×10 min, 45°C; 1.5X SSC buffer, 1×10min, 45°C; 2x SSC buffer, 2×20 minutes, 37°C. After washing steps, sections were incubated with 1X blocking solution (Biochain) in PBS for 1 hour. Slides were incubated overnight with alkaline-phosphate conjugated anti-digoxinogen antibody (diluted 1:500 in PBS) (Biochain) at 4°C. The following day, slides were washes three times with PBS for 10 minutes, twice with alkaline phosphatase buffer (Biochain) for 5 minutes, and incubated with nitro-blue tetrazolium chloride (NBT) and 5-bromo-4-chloro-3’-indolyphosphate p-toluidine salt (BCIP) solution (6.6 µL NBT and 3.3 µL BCIP were diluted in 1 mL alkaline phosphatase buffer) (Biochain) for 20 hours at room temperature. Slides were scanned using a ScanScope CS system (Aperio; Vista, CA).

### Assays

Insulin was measured in plasma, islets and cell culture media using a Mouse Ultrasensitive Insulin ELISA (ALPCO, Salem, NH). Proinsulin was measured from plasma collected from cardiac blood using Rat/Mouse Proinsulin ELISA (Mercodia, Sweden). To perform migration assay with cultured cell lines, PANC-1 cells were transfected with the miR-216a and control scrambled miRNA mimetics (Dharmacon, Lafayette, CO) and 24 hours later, seeded on trans-well migration chambers (Trevigen, Gaithersburg, MD) in media without FBS. Cells were allowed to migrate for 16 hours and the number of cells transversing the boyer chamber was quantified by incubating the cells in the bottom chamber with Calcein-AM for one hour and measuring the fluorescence at 485 nm excitation, 520 nm emission with a Tecan Plate Reader. For the XTT (2,3-bis-(2-methoxy-4-nitro-5-sulfophenyl)-2H-tetrazolium-5-carboxanilide) assay, cells were transfected with miR-216a and the control miRNAs and an XTT assay was performed 3 days post-transfection in 96-well plates. XTT (Life Technologies) was dissolved in pre-warmed 37°C cell culture media at 1 mg/mL and stock Phenazine methosulfate (PMS) solution was prepared in PBS at 10 mM. PMS was mixed with the XTT solution immediately before labeling the cells and 25 µL of XTT/PMS solution was directly added to each well containing 100 μL cell culture media. Cells were incubated for two hours at 37°C in a CO_2_ incubator. Absorbance was read at 450 nm using a Tecan Plate Reader. For live cell imaging, INS-1 cells seeded on 96-well plates were transfected with control or miR-216a expressing plasmids or left untreated. Two days after transfection, cells were incubated with 50 ng/mL Hoechst and 1:500 diluted Alexa647 annexinV 30 min prior to imaging. TNF-α, IFN-ϒ and IL-1β (10 ng/mL each) was added to media and cells were imaged every 2 hours at 37°C and 5% CO_2_ in an ImageXpress Micro™ (Molecular Devices).

### RNA Sequencing and Analysis

RNA sequencing was performed by the Biomedical Research Center Genomics facility at the University of British Columbia, Vancouver, Canada. Sample quality control was assessed using an Agilent 2100 Bioanalyzer. Qualifying samples (samples with RNA integrity numbers > 8) were then prepared following the standard protocol for the NEBnext Ultra ii Stranded mRNA (New England Biolabs). Sequencing was performed on the Illumina NextSeq 500 with Paired End 42bp × 42bp reads. Sequencing data were demultiplexed using Illumina’s bcl2fastq2.

Fastq files were retrieved from Illumina Basespace and aligned with salmon^65^ (version 0.13.1) to the most current Ensembl Mus Musculus genome (as of 2019-05-08)^66^ using the gcBias, validateMappings, and rangeFactorizationBins (with 4 as the binning parameter) options. Quality of the Fastq files was assessed using FastQC^67^ (version 0.11.8). All fastq files passed quality control. All further analyses were performed in R (version 3.6.0)^68^. Counts were imported using the tximport package (version 1.12.0)^69^ and the biomaRt package (version 0.8.0)^70,71^. Analysis of differentially expressed genes was performed using the DESeq2 (version 1.24.0)^72^. Genes with less than 5 counts in 2 samples or fewer were dropped from the analysis. Gene Ontology analysis was performed on genes with an IQR greater than 0.5 across all samples and an adjusted p value < 0.05 (adjusted by Benjamini-Hochberg method) between wild type and control mice. Significantly enriched terms for Biological Process, Cellular Components, and Molecular Functions were identified using the categoryCompare package (version 1.28.0)^73^ and visualized using Cytoscape via the RCy3 package (version 2.4.0)^74^. Further KEGG network analyses were performed using the gage (version 2.34.0)^75^ and pathview (version 1.24.0) packages^76^. Finally, GO terms were summarized using the REViGO methodology^77^ and plotted using the treemap (version 2.4.2) package^78^. All R scripts and data are available upon request.

### Statistical Analyses

Statistical analyses were performed using GraphPad Prism 7, with significance defined as p < 0.05. Unless specified in the figure legend, student’s t-tests were used for normally distributed data (two-tailed, unpaired). Bonferroni correction was used to correct for multiple comparisons, where applicable. Data are represented by bar plots with individual values indicated, individual values only, or line graphs. Results show mean±SEM.

## Acknowledgements

We thank Dr. Anna D’Souza for providing islets from Leptin knock-out rats. T.J.K. gratefully acknowledges funding from JDRF and the Canadian Institutes for Health Research for this research. S.E. is a recipient of a JDRF advanced postdoctoral fellowship. We would also like to thank Dr. Ziliang Ao and Dr. Garth L. Warnock from the Irving K. Barber Human Islet Isolation Laboratory (Vancouver, BC) for providing human islets.

---

**Suppl. Figure 1.**
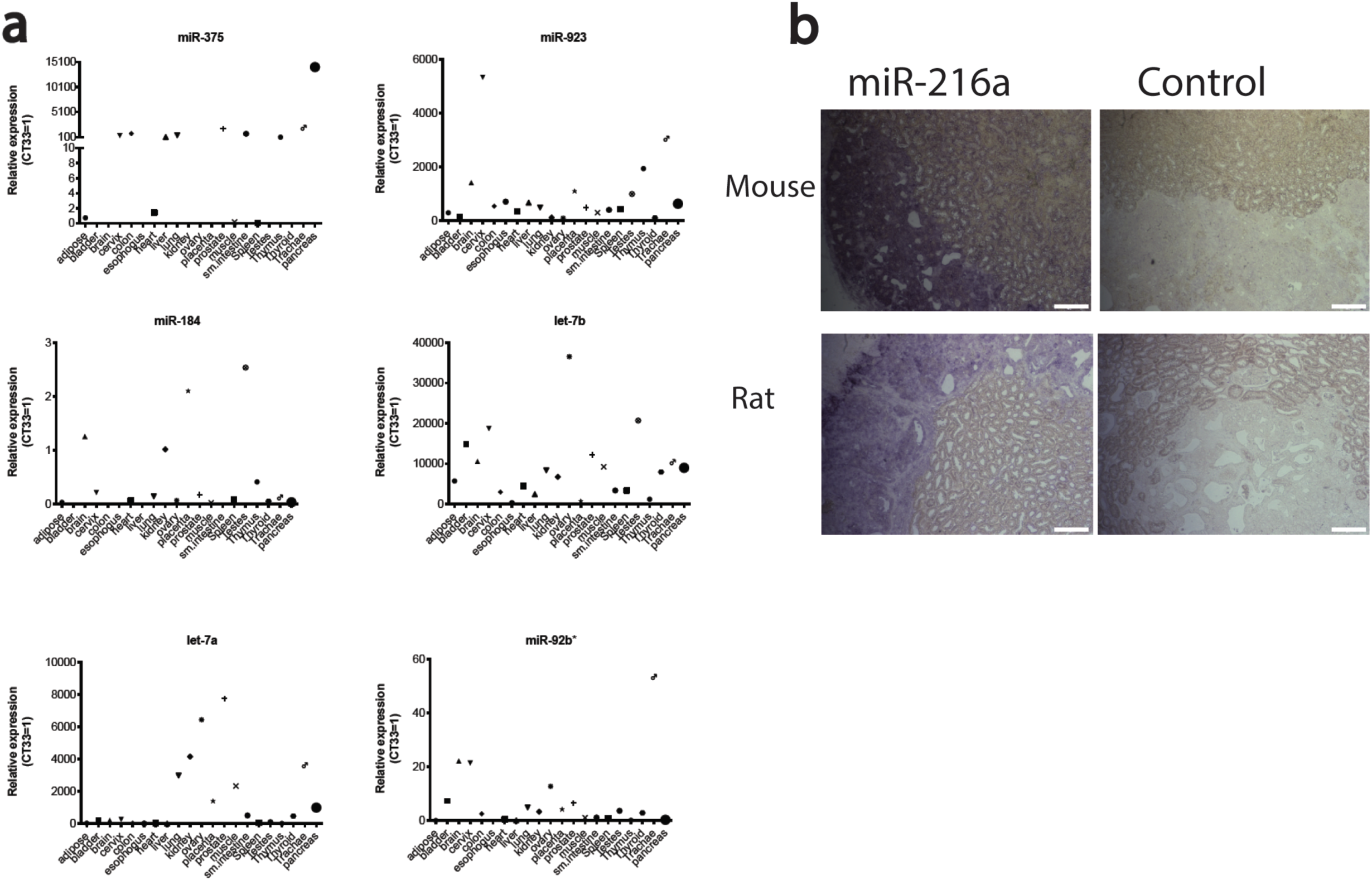
**a** Expression of the indicated miRNAs in various human tissu es. Equal amounts of RNA from human tissues (each a pool of 3 tissue donors) was reverse-transcribed and expression of indicated miRNAs was determined by qRT-PCR. Threshold cycle 33 (Ct= 33) was arbitrarily set as 1. **b** miR-216a is expressed in pancreatic tissue differentiated from hESCs. Representative in situ hydridization images of differentiated hESC-derived grafts at 22 weeks post implant in a mouse and a rat. Grafts harvested from mice and rats were probed with DIG-labeled miR-216a and scrambled miRNA control probes. Scale bars= 100 µm.

**Suppl. Figure 2.**
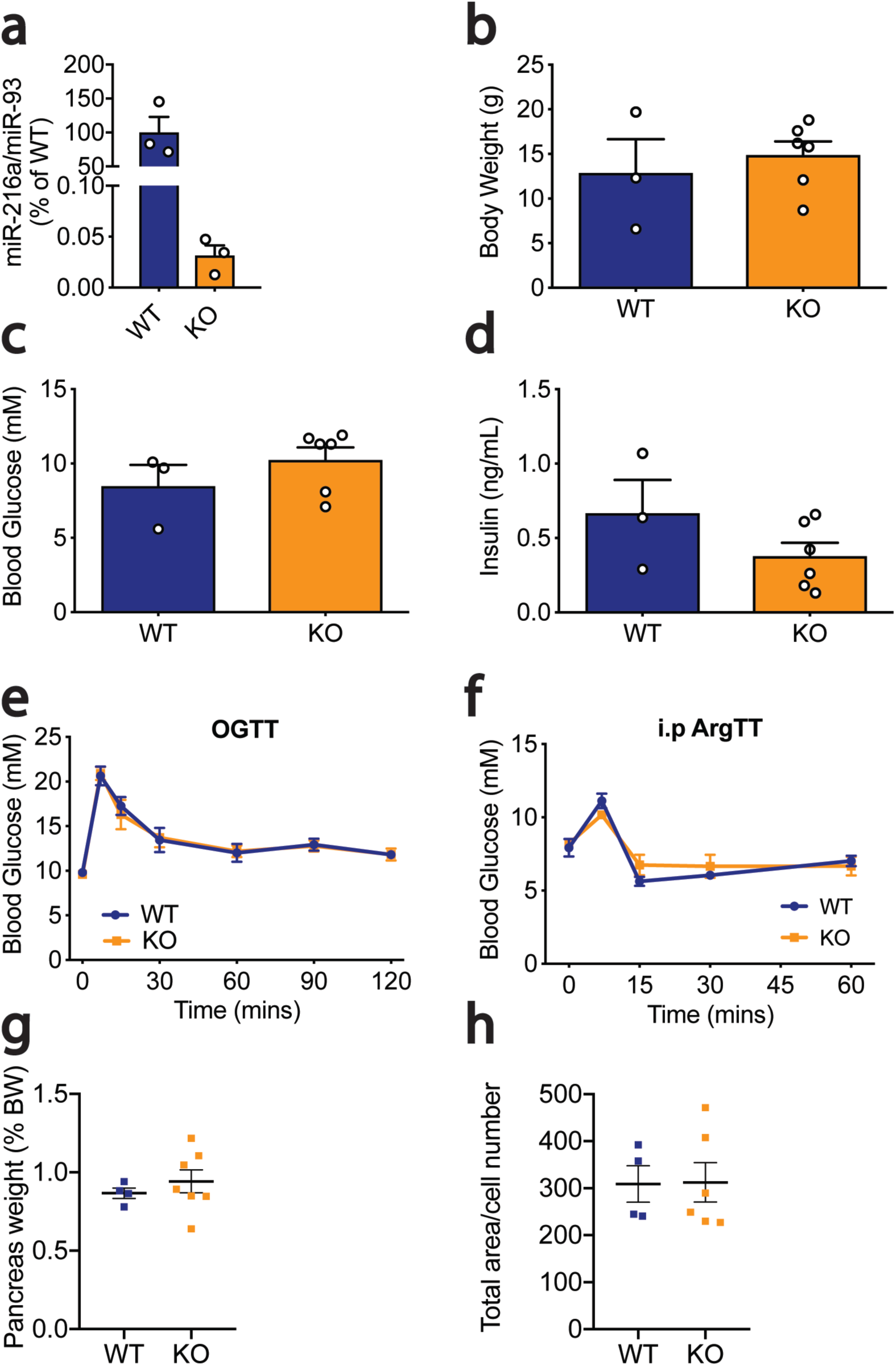
miR-216a KO mice do not display alterations in glucose homeostasis. **a** Islets from 10-week old male WT and miR-216a KO mice were isolated and miRNA expression was quantified with qRT-PCR. n = 3. **b-d** Body weight **(b)**, fasting blood glucose **(c)**, and insulin measurements **(d)** from 3-4 week old male mice. n = 3-6. n.s. =non-significant.A two-tailed Student’s t-test was performed to assess significance. **e** Oral glucose tolerance (OGTT) test performed on 15-week old mice. **f** Intraperitoneal arginine tolerance test (i.p. ArgTT) performed on 18-week old mice. n = 4-6. **g** Pancreata from 21-week old WT and miR-216a KO mice were weighed and normal ized to body weight. **h** Pancreatic cell size was assessed by analyzing dapi staining from the pancreata of 21-week old male mice. Individual data points are shown in **(a-d, g-h).** Data represent mean± SEM.

**Suppl. Figure 3.**
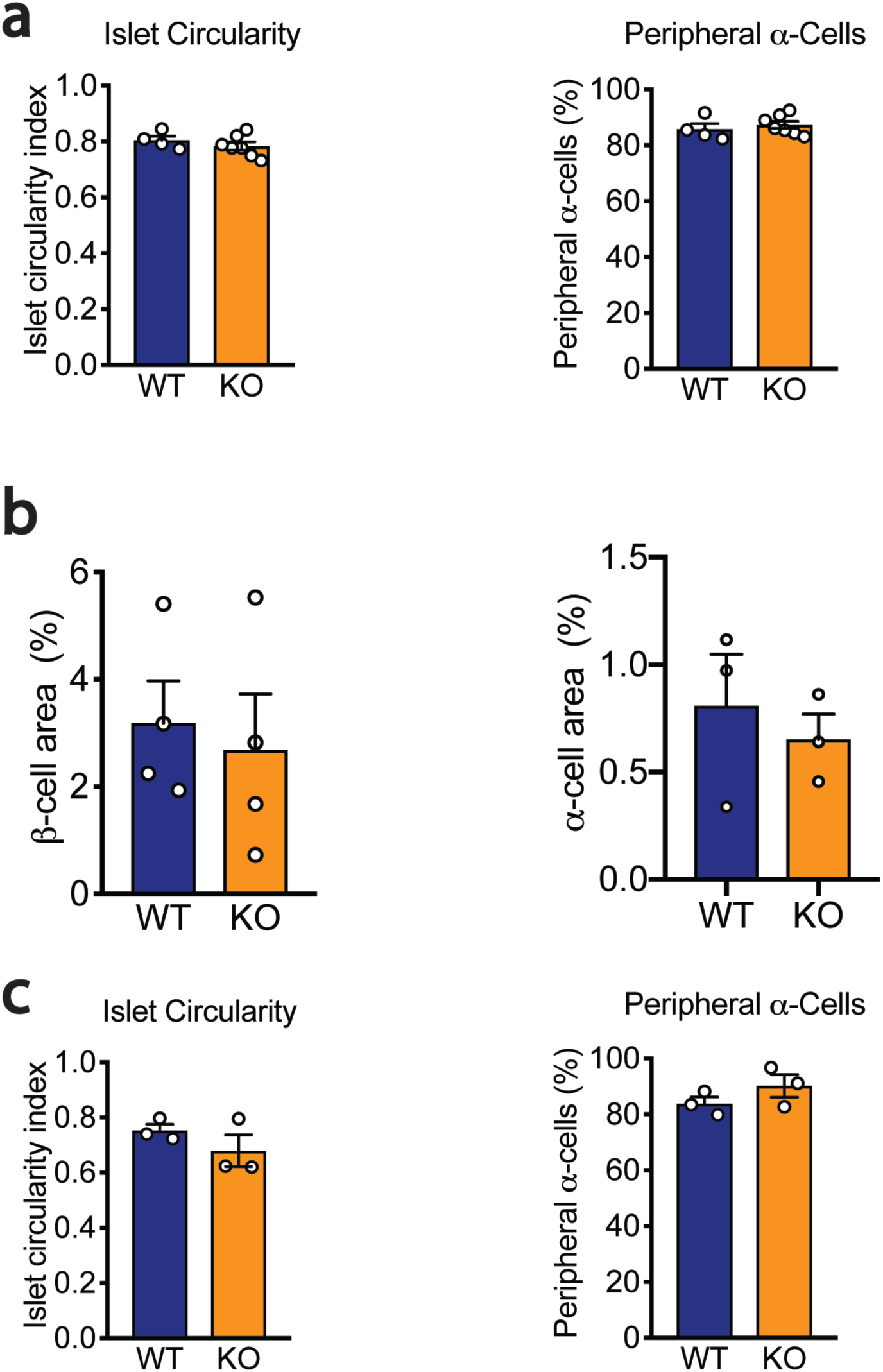
Islet circularity and peripheral α-cell numbers are unchanged in miR-216a KO mice. Pancreata from adult male mice **(a;** n=4-7) and 1-day old male neonatal mice **(b-c;** n = 3-4) were immunostained for synaptophy sin, insulin and glucagon, and islet circularity, peripheral α-cell percentage, β-cell area, and β-cell area were calculated. Individual data points are shown. Data represent mean ± SEM.

**Suppl. Figure 4.**
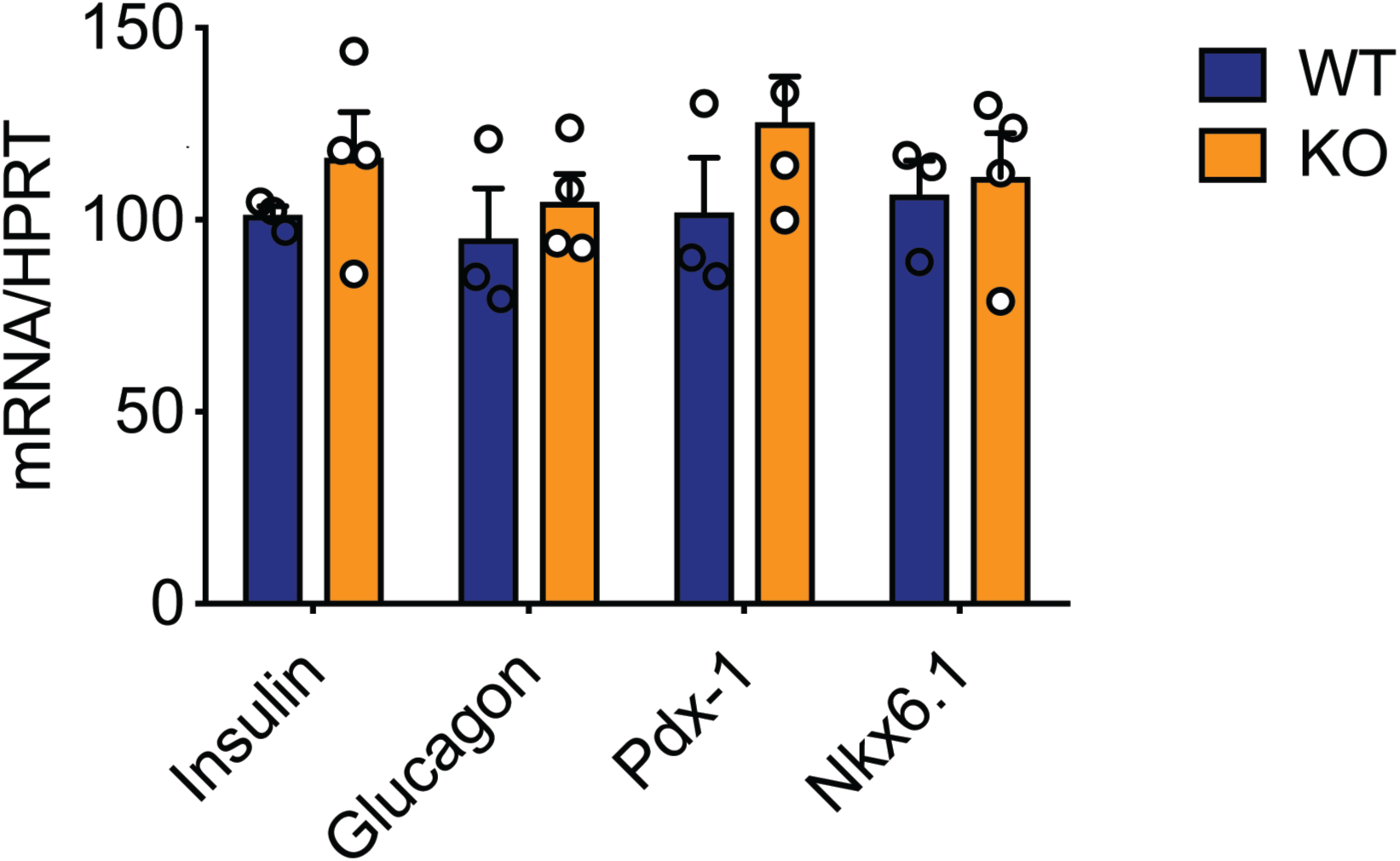
Gene expression analysis using islets isolated from 10-week old male WT and miR-216a KO mice. RNA was isolated, reverse-transcribed and expression of the indicated genes was determined by qRT-PCR. WT levels arbitrarily set as 100. n = 3-4. Individual data points are shown. Data represent mean ± SEM.

**Suppl. Figure 5.**
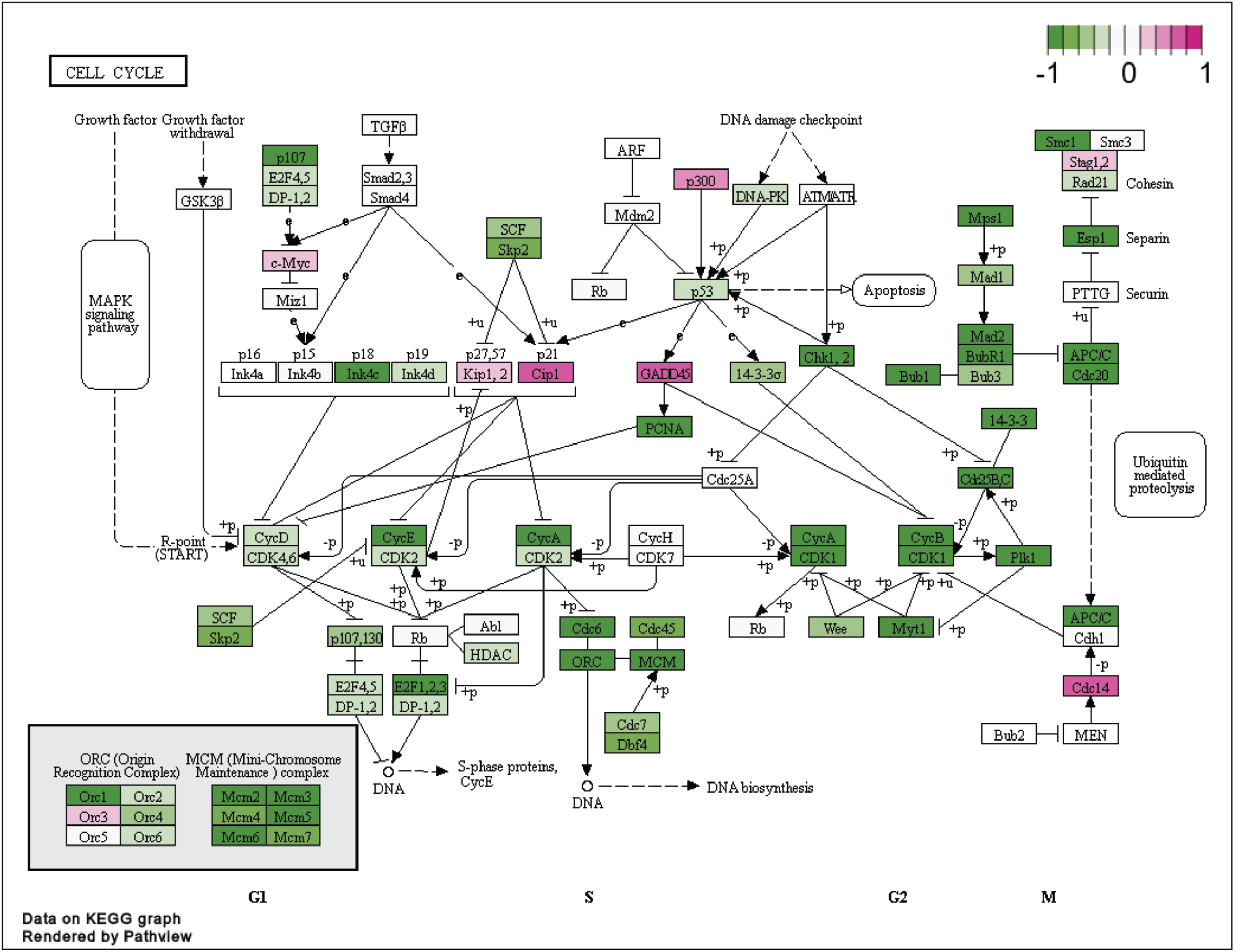
**A cell cycle pathway map,** with genes significantly upregulated in miR-216a KO mice compared to WT mice shown in purple, and genes significantly downregulated in the KO mice compared to WT mice shown in green. Cell cycle is a significantly enriched GO “Biological Process” and KEGG term, with statistical significance defined as a q value < 0.1 (therefore allowing a 10% FDR).

